# On-skin, micro-objective enabled camera module for speckle contrast optical spectroscopy/tomography

**DOI:** 10.1101/2025.06.12.659250

**Authors:** Andres Quiroga, Lorenzo Cortese, Manish Verma, Peter Dannberg, Ilias Tachtsidis, Norbert Danz, Turgut Durduran

## Abstract

In this paper, we introduce a speckle contrast optical spectroscopy/tomography (SCOS/SCOT) configuration based on an integrated imaging approach exploiting 113 micro-objetives mounted on a commercial CMOS camera that operates without fiber coupling, suitable for direct skin contact measurements and simultaneous multiple source-detector separation acquisitions. This compact system was validated *ex vivo* on phantoms and *in vivo* by monitoring the blood flow on the forearm muscle of a healthy human subject. The measurements, performed at multiple source-detector separations and camera exposure times, demonstrate excellent agreement with the theory based on the correlation-diffusion model. *In vivo* data demonstrate the capability of tracking pulsatile blood flow with a high signal-to-noise ratio (*>*4 harmonics of the cardiac pulse frequency detected) and sensitivity to small changes in muscle blood flow. This micro-objective array-based design overcomes a key challenge towards wearable SCOS/SCOT devices.

## 1. Introduction

Speckle Contrast Optical Spectroscopy and Tomography (SCOS/SCOT) [1–5] is an emerging modality for measuring and imaging local, microvascular blood flow in tissue using diffuse laser speckle statistics in the near-infrared. Typically, SCOS/SCOT measurements^1^ utilize source-detector geometries that enable measurements deep (*>* 1 cm) inside the tissues. SCOS offers an advantageous alternative to the better established diffuse correlation spectroscopy (DCS) that also is based on the same speckle statistics, in the sense that SCOS is able to work with less constraints in timing (∼ 100s of nano-seconds versus∼ milliseconds) and can readily be massively parallelized (10^5^ or more independent speckles). The physics of SCOS allows for signal acquisition through temporal integration over relatively long detector exposure times. This capability enables the use of widely available, industrial-grade CMOS cameras—featuring high pixel counts and low cost—rather than relying on expensive, specialized high-speed detectors [6, 7].

Originally, SCOS was demonstrated as a non-contact system [1–4] utilizing common objective lenses with a careful arrangement of the system aperture with a *>* 10 cm working distance which is not a robust way to measure living tissues due to subject motion, discomfort, stray light influences and others. This approach was largely abandoned (except for specific applications such as surgical flap characterization) [8], and was replaced multimode fibers coupled directly to the cameras [5, 9]. A common configuration uses a multimode fiber (e.g. 200 to 800 *μ*m core) to collect scattered light, which is then projected by the micro-objective onto a portion of large-format CMOS sensor.

However, ideally, one would like to abandon the uses of fibers whose modal-noise and subtle vibrations add noise to the measurements, whose bulk and fragility limit the applications, and, the number of parallel cameras/fibers limit the spatial sampling. Several attempts were made to place the detector arrays directly on the tissues such as those utilizing single-photon avalanche photo-diode arrays [10, 11] with a ninety-degree objective/prism combination, bulky camera/objective combinations with complex thermal management [12] and with low signal-to-noise-ratio (SNR) minimal foot-print wafer-level cameras [13]. These attempts all face the issue that the lack of appropriate lens/aperture combinations meant either a bulky, impractical applications or limited SNR. We have hypothesized that the wide-spread, replicated micro-optical approaches could be adapted for the needs of on-skin, wearable SCOS detection which would then pave the way for the freedom to choose the appropriate cameras. These designs would further enable new approaches such as single-camera multi-distance, highly parallelized measurements.

To that end, here we introduce a SCOS configuration based on a micro-objective-enhanced CMOS camera that operates without fiber coupling [14, 15]. This approach allows the camera to be placed directly in contact with the skin, eliminating the need for fiber optics, maximizing use of the entire sensor surface and enabling simultaneous multi-distance acquisitions. A micro-objective array mounted in front of the sensor segments the incoming speckle field into multiple independent sampling regions (i.e. lens field of view), each corresponding to a distinct source-detector geometry.

This configuration offers several advantages. First, it enables simultaneous acquisition of multiple speckle patterns and multi-distance source-detector detection, allowing for estimation of blood flow at different positions. Second, it allows multi-exposure acquisition, expanding the dynamic range for detecting both fast and slow flow regimes. Finally, fiber elimination results in a compact and lightweight probe suitable for handheld or portable implementations.

In this study, we present a proof-of-concept demonstration of this system with a generic CMOS camera. We characterize its performance on static and dynamic phantoms, and validate its ability to capture reliable speckle patterns from human skin, estimate the speckle contrast, and extract hemodynamic information. This represents a significant step toward compact, scalable, and high-throughput SCOS devices for clinical and research applications.

## 2. Methods

### 2.1. Experimental setup

The SCOS system consisted of three primary components: (i) a fiber-coupled coherent illumination source, (ii) a micro-objective-enhanced miniaturized CMOS imaging module, and (iii) a modular sample interface (probe). The light source consisted of a single longitudinal mode laser emitting at 785 *nm* (iBEAM SMART 785, Toptica, Germany), fiber coupled to a three-meter multimode fiber of 200 *μm* core (FT200EMT, Thorlabs, US).

The custom miniaturized imaging module [14, 16] was developed at Fraunhofer Institute to enable large area microscopic imaging with an integrated, compact system. The optical design of the system is depicted in Fig. 1 and consists of a hexagonally arranged array (Fig. 1 b) of identical micro-optical microscopes, each yielding approximately 4.8x magnification and NA 0.24 observation. The system consists of four major elements: the array of micro-objective, a long pass filter to avoid short wavelength illuminating the sensor (LP605ET, Semrock, US), an anti-cross talk element to suppress light exchange in-between neighboring channels, and an image sensor. The latter is a commercial CMOS camera (VCXU-201M.R, Baumer, Germany) with Sony IMX 183 image sensor (active area 5472 × 3648 pixels, pixel size 2.4 *μm*).

**Fig. 1.**
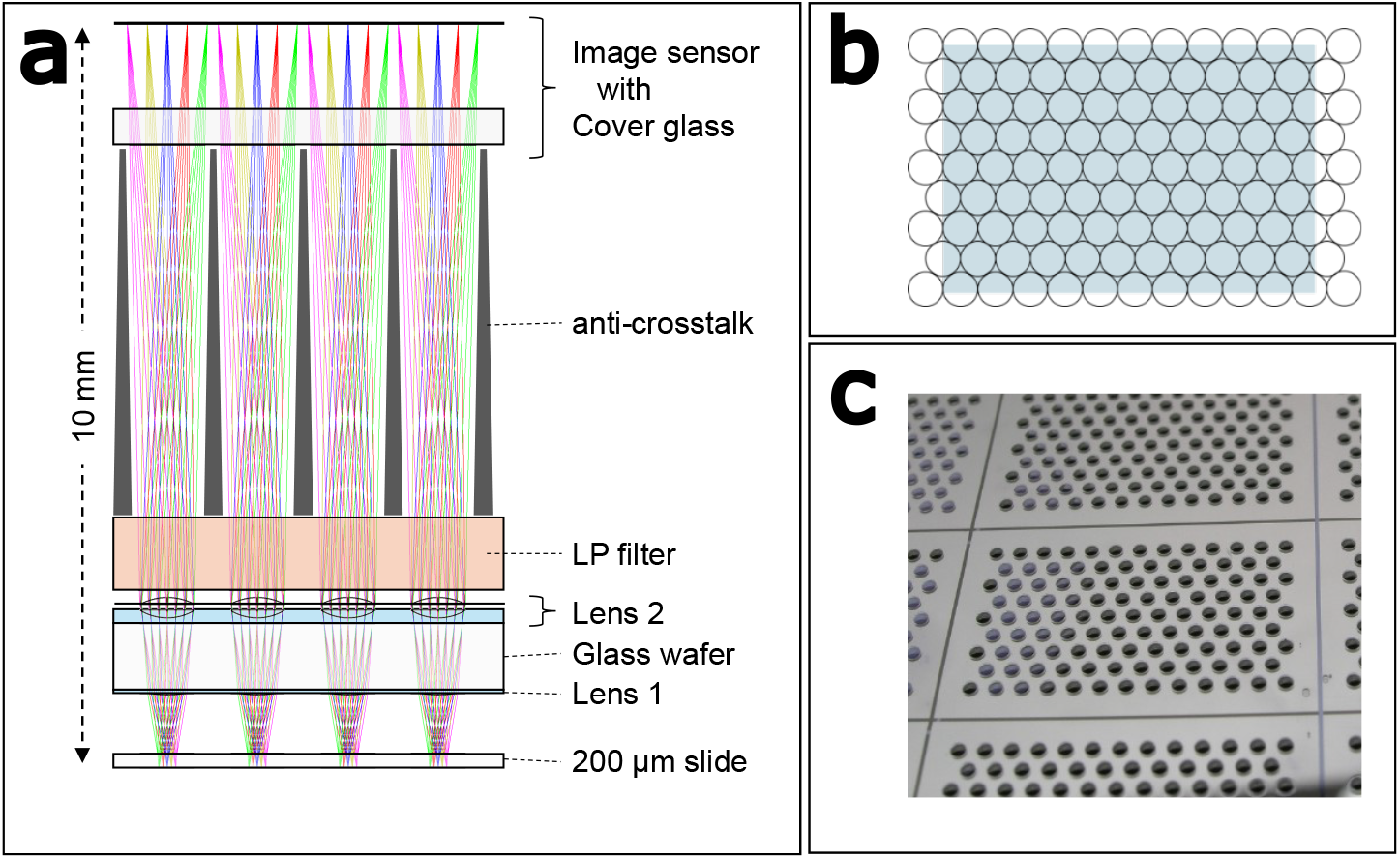
(a) Optical raytracing layout of four channels illustrating the array microscope set-up. Lens 1 is hardly reproduced in this sketch due to it low sag-height; aspherical lens 1 and achromatic lens 2 have been prepared by polymer-on-glass replication. (b) Hexagonal arrangement of the array channels relative to the image sensor indicate light blue. (c) Photograph of the replicated objective array prior to wafer dicing.

Preparation of the micro-optics follows the procedures described previously [17]. Master structures for replicating the micro-optical elements have been prepared either by reflow technology (spherical surfaces of the achromatic lens 2 in Fig. 1) or by ultra precision machining (aspherical lens 1, UPT, Nuremberg, Germany). Copies of these structures have prepared by thermal crosslinking an elastomeric layer on glass to be used as replication tools. Prior to polymer replication, black apertures have been prepared on both sides of a glass wafer by lithographic patterning of spin coated black matrix polymer. In a last step, lenses have been aligned and replicated onto the aperture structure in a mask aligner (SUSS MA-8) using UV curable resins, where the achromatic lens requires two successive replication steps using different polymers.

The anti-crosstalk element has been mechanically processed from Aluminum with subsequent anodization to suppress reflection of the side walls. It is worth noting that it combines three different functions. First, it is the spacer to ensure the correct distance between image sensor and optics, second it denies inter-channel optical cross talk, and last not least it is the mechanical holder for inserting filter and optics, thus enabling a very simple system integration without any active alignment step.

Prior to integration, the camera’s housing must be revised to remove the standard adapter, to access the image sensor directly and to enable the object to get close enough to the system. In conclusion, each channel in the array features a field-of-view of approximately 230 *μm* in diameter and yields an image size of approximately 1100 *μm* in diameter with a pitch of 1250 *μm*.

**Table 1.** Baumer VCXU-201M.R Camera Specifications.

| Feature | Specification |
| --- | --- |
| **Model** | VCXU-201M.R |
| **Manufacturer** | Baumer |
| **Sensor** | CMOS Sony IMX183 |
| **Camera type** | Monochrome |
| **Resolution** | $5472 \times 3648$ (20 MP) |
| **Sensor Size** | 1-inch ($11.264\text{ mm} \times 5.984\text{ mm}$) |
| **Pixel Size** | $2.4\text{ }\mu\text{m} \times 2.4\text{ }\mu\text{m}$ |
| **Pixel Format** | Mono8, Mono10, Mono12, Mono12p |
| **Frame Rate** | 15 fps @ Full Resolution |
| **Interface** | USB 3.0 (5 Gbps) |
| **Shutter Type** | Global Shutter |
| **Bit Depth** | 8 / 10 / 12-bit |
| **Relative Response** | 0.5 (785 nm) |
| **Operating Temperature** | 0°C to 65°C |
| **Dimensions** | $29\text{ mm} \times 29\text{ mm} \times 47\text{ mm}$ |
| **Weight** | 90 g |

The laser source fiber and the imaging module were then fixed on a custom built probe (see Figure 2) which consisted of a plastic cylindrical support (diameter 5 *cm*) for keeping the probe by hand (as for the *in vivo* experiments) and on fixed supports (as in the phantom experiments), and a passive cooling system, to keep the camera temperature at the optimal operating temperature of∼ 45 °C, while keeping the plastic cylindrical support below ∼ 37 °C, suitable for skin contact. The micro-objective array was covered by a glass slide of 200 *μm* thickness, with the outer surface placed at the working distance (900 *μm*) from the lens system, enabling direct skin contact measurements. This probe configuration allowed simultaneous detection of signals from multiple source-detector separations, starting from 11 *mm*. As reported in the following sections, acquisitions have been performed on a limited region of interest (ROI) of the camera sensor, limiting the source-detector separations available to the range between ∼ 11 and ∼ 23 *mm*.

**Fig. 2.**
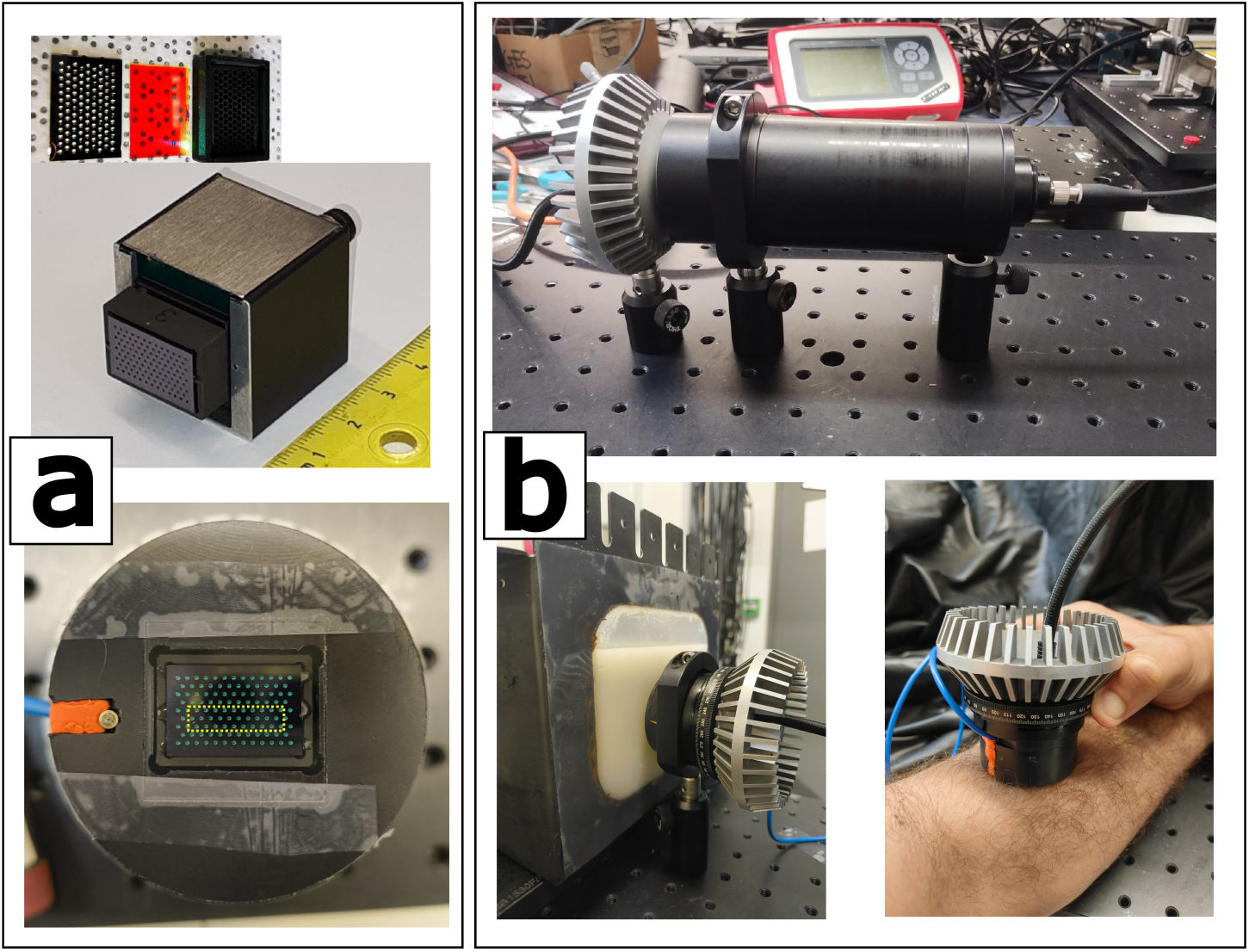
(a) Photo of the miniaturized imaging module and its components (top panel), and mounted into the probe (bottom panel, in dashed line we have highlighted the ROI used in liquid phantom and *in vivo* measurements). (b) Experimental configurations for static phantom (top), liquid phantom and *in vivo* experiments.

### 2.2. Experimental Protocol

Three different validation experiments were conducted: (i) validation on a static phantom; (ii) on a dynamic phantom; (iii) *in vivo*.

#### Static phantom experiment

As a first characterization of the SCOS system, we have analyzed the speckle pattern generated by illuminating laser light on a glass diffuser (DG20-1500, Thorlabs, US), and detected in transmission geometry. This measurement allowed the determination of the speckle size, the number of the speckles within the field of view of each lens, and the system noise. This experimental setup was enclosed in a black tubing to reduce ambient light contamination. The glass diffuser was placed in contact with the probe (i.e., at the working distance of the micro-objective array) and laser power was tuned in order to optimize the signal detected in the camera. The measurement consisted of 60 *s* acquisition at a rate of 20 *Hz* and camera exposure time *T* = 3 *ms*. Before the measurement, background signal was acquired with no laser illumination.

#### Dynamic phantom experiment

*Ex vivo* tests were performed on a tissue simulating liquid phantom consisting of a suspension of fat droplets (Lipofundin 20%, B. Braun Melsungen AG, Germany) in water, with components’ concentrations chosen to obtain a reduced scattering coefficient 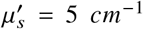 and absorption *μ*_*a*_ = 0.026 *cm*^−1^ [18]. The probe was fixed on a support and placed in contact with the transparent window of the phantom container (see Figure 2). Phantom and probe were then covered with a black cover to reduce ambient light contamination and laser power was set to 50 *mW* at the fiber tip to avoid saturation effects in the camera. Each measurement consisted of sixty-second acquisitions at 20 *Hz* acquisition rate.

Measurements were repeated at different camera exposure time (*T*) ranging from 1 *ms* to 5 *ms*. Each measurement was preceded by the acquisition of the dark signal at the same exposure time. In order to reduce the amount of data and to improve the acquisition rate up to 20 *Hz*, each acquisition was limited to the region of interest (ROI) represented in Figure 2, allowing source-detection separations ranging from ∼ 11 to 23 *mm*.

#### *Iv vivo* experiment

As a proof of concept for validating the capability of the system to track pulsatile blood flow, we have monitored the blood flow in the forearm muscle of a healthy human subject for one minute. This experiment was approved by the local Ethical Committee (ref. ICFO_HCP/2012/1) and was conducted according to the guidelines of the Declaration of Helsinki. The subject signed an informed consent form prior to participation. The hand-held probe was placed on the subject’s forearm muscle (*brachioradialis*) and held steady by the operator’s hand (see Figure 2), minimizing motion artifacts. The laser power injected into the tissue was set to 90 *mW* and pulsed at 20 *Hz* (synchronized with the camera acquisition rate), with a pulse duration of 10 *ms*, in accordance with the maximum permissible exposure (MPE) limits for skin safety defined by the International Organization for Standardization (ISO) and the American National Standards Institute (ANSI). Measurements were repeated at different camera exposure times, ranging from *T* = 2 *ms* to *T* = 5 *ms*. As in the phantom experiment, each measurement was preceded by background signal acquisition, and each acquisition was limited to the ROI shown in Figure 2.

### 2.3. Data Analysis

Raw images were acquired at 20 *Hz* in 16-bit format and were shifted by 4 LSBs (Least Significant Bit) to match the camera 12-bit resolution. Gray values in Digital Numbers were converted to electrons using the manufacturer’s gain calibration, defined as the ratio between full-well capacity (14959 electrons) and the 12-bit maximum count (4095 ADU). The resulting electron counts were corrected by subtracting the mean dark current obtained from a series of dark frames.

To define analysis regions for *κ*^2^ quantification, binary masks based on the micro-objective array were applied. In each lens field of view we have selected a circular region with a radius ∼ 60% of the lens radius (approximately 17180 pixels per circle, as example, see Fig. 3 bottom left panel, and Fig. 4 top panel) to reduce edge artifacts and ensure uniform illumination.

**Fig. 3.**
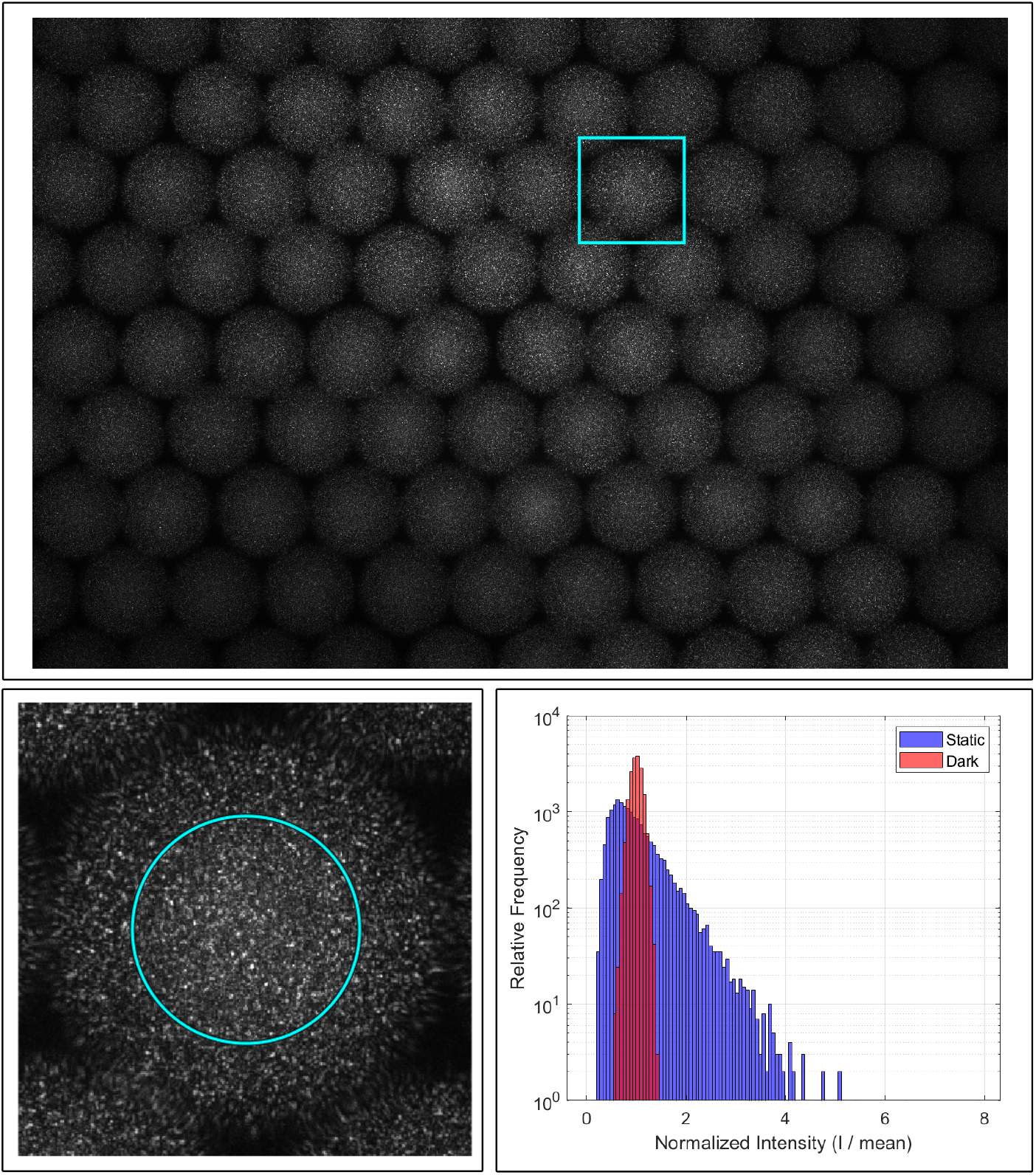
Static phantom experiment. Top: Image of the whole sensor. Bottom left: Zoomed image of the field of view of one micro-objective. Bottom right: Intensity distribution histogram from the selected lens field of view, comparing dark distribution and intensity distribution obtained from the static phantom.

**Fig. 4.**
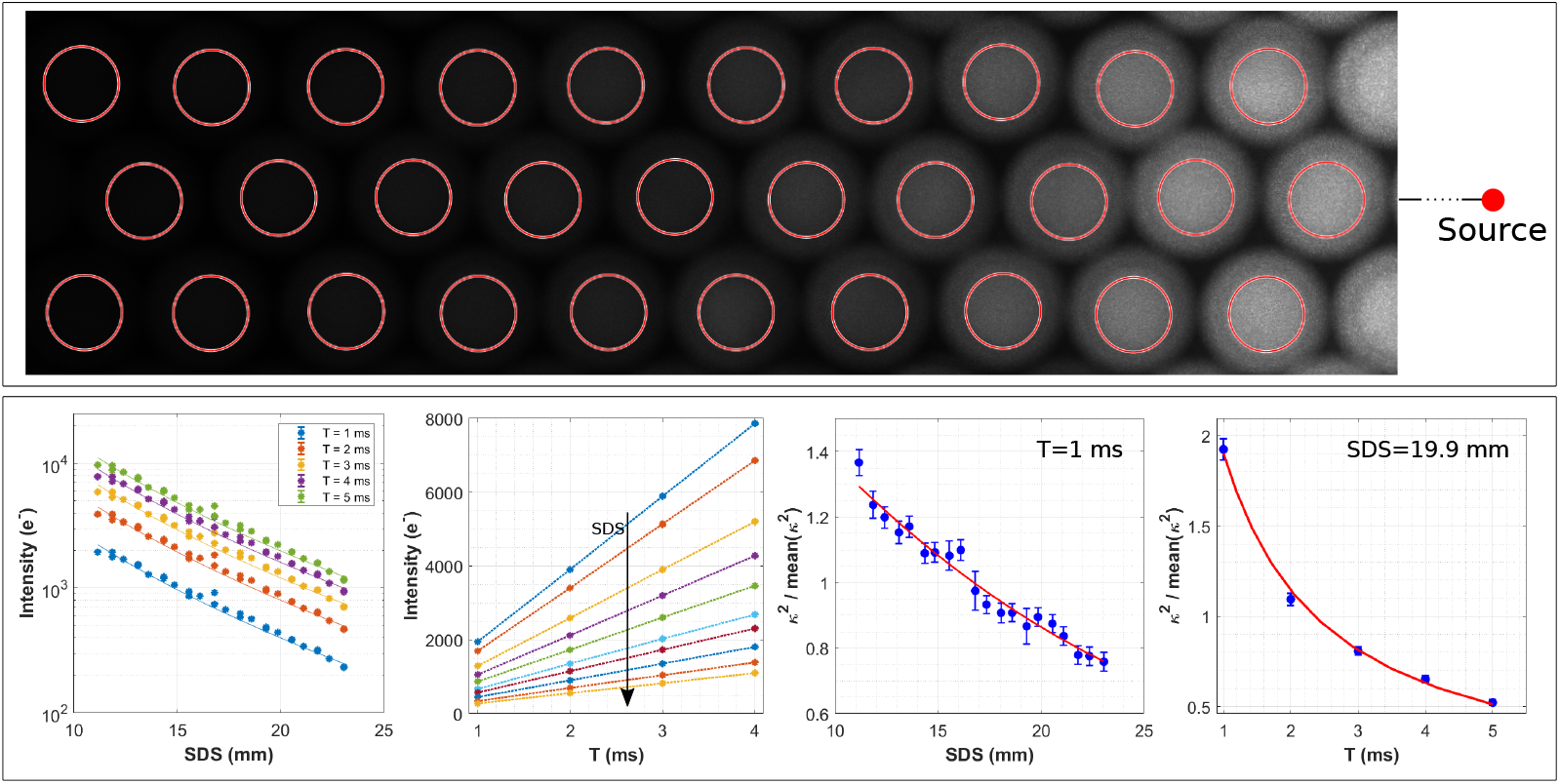
Liquid phantom experiment. Top panel: Exemplary image frame of the ROI (*T* = 3 *ms*). Bottom panel: Intensity *I* and speckle contrast *κ*^2^ Vs. source-detector separation (SDS) and exposure time *T* together with theory expectations (bulk lines). Source-detector separations range from 11.15 *mm* to 23.05 *mm*.

The speckle contrast, *κ*^2^, was obtained from the raw intensity statistics of each frame within each lens field of view. Following the approach described in [1], *κ*^2^ was computed by correcting the spatial variance for both photon shot noise and camera dark noise, using the expression:

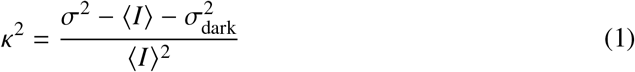

where *σ*^2^ is the spatial variance of the intensity in the field of view, ⟨*I*⟩ is the mean intensity, and 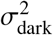 represents the variance estimated from dark frames acquired with the illumination source off. Speckle contrast signals were subsequently inverted to compute the Blood Flow Index (BFI) as 1 /*κ*^2^ (*T*).

The speckle contrast depends on the electric field correlation function through the relation [1]:

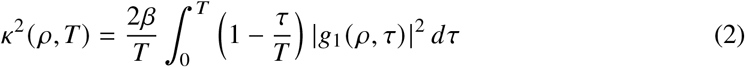

where *g*_1_ is the normalized electric field autocorrelation function, and *β* is an experimental constant mainly depending on the optical system.

To compare speckle contrast dependence on source detector separation and exposure time with theory in the dynamic phantom experiment, we have determined the Brownian diffusion coefficient of the particles in suspension by fitting the experimental data, for varying source-detector separation or exposure time, with equation 2. In the fit, *g*_1_ was calculated as the solution of the correlation diffusion equation for a semi-infinite homogeneous medium in reflectance geometry [1, 19]. Since the experimental parameter *β* was not measured, experimental and theoretical speckle contrasts have been normalized by its mean.

Additionally, the dependence of the measured intensity on source-detector separation and exposure time was then verified by using Beer-Lambert law for a semi-infinite homogeneous medium geometry [1, 20].

Lastly, to quantify the capability of the system to track pulsatile flow *in vivo*, we have calculated the Fourier spectral amplitude of the speckle contrast related to the one-minute measurement on the forearm muscle at multiple source-detector separations and exposure times, together with the signal-to-noise ratio SNR. The SNR was calculated as the ratio of peak power in the heartbeat frequency band (0.8-1.2 Hz) to the mean power of the baseline noise, estimated between the first and the second harmonic of the heartbeat frequency [21–23].

## 3. Results

### Static phantom experiment

In Figure 3, we present an exemplary acquisition in transmission geometry from the static phantom (i.e., a glass diffuser), including the full active area of the sensor (113 micro-objectives, top panel), a representative signal from the field of view of a single micro-objective (bottom left), and the corresponding intensity histogram comparing the dark distribution to the diffuser signal. As discussed in Section 2.3, the effective area for speckle contrast analysis was limited to 60% of the micro-objective radius, yielding a circular mask with a radius of 73.95 pixels in the binned image (binning 2). This corresponds to a total of 17180 pixels within the mask.

The speckle size was independently estimated from unbinned (binning 1) images using spatial autocorrelation analysis, yielding an average speckle diameter of 3 pixels (∼ 7.2 *μ*m). To match the binned image (pixel size doubled), this would correspond to 1.5 pixels in binning 2. Based on this, the estimated number of speckles within the mask is approximately 5469, corresponding to an average of about 3.14 pixels per speckle.

### Dynamic phantom experiment

In Figure 4 (top panel) we report a representative frame of the ROI acquired with sensor exposure time *T* = 5 *ms*. The image shows an attenuation of the intensity of the speckles as source detection separation increases, as expected. This effect is quantified and reported in Figure 4 (bottom panel), where the average intensity of the image related to each micro-objective field of view is reported for varying source-detector separations and exposure times together with theory expectations. In the same figure, the dependence of the speckle contrast *k*^2^ is also reported for varying source-detector separation (at a fixed exposure time *T* = 1 *ms*) and exposure time (at a fixed source-detector separation *SDS* = 19.9 *mm*), and compared with the theoretical model (equation 2).

### *Iv vivo* experiment

In Figure 5 we report an exemplary image frame of the ROI (exposure time *T* = 3 *ms*) acquired in the forearm muscle of a healthy subject, together with the *BF I* pulsatile signal obtained from the speckles imaged at a frame rate of 20 *Hz* by each of the lenses of the ROI (zoom of five-seconds time interval over the complete measurement of one minute). Additionally we report the Fourier transform of the *BF I* obtained from the lens #20 in a time window of one minute, highlighting the oscillations of the signal due to the subject heartbeat frequency (∼ 1 *Hz*). Fourier analysis furthermore highlighted high signal-to-noise ratio (SNR>250 and >4 harmonics visible) for all the source-detector separations and exposure times (data not shown) utilized.

**Fig. 5.**
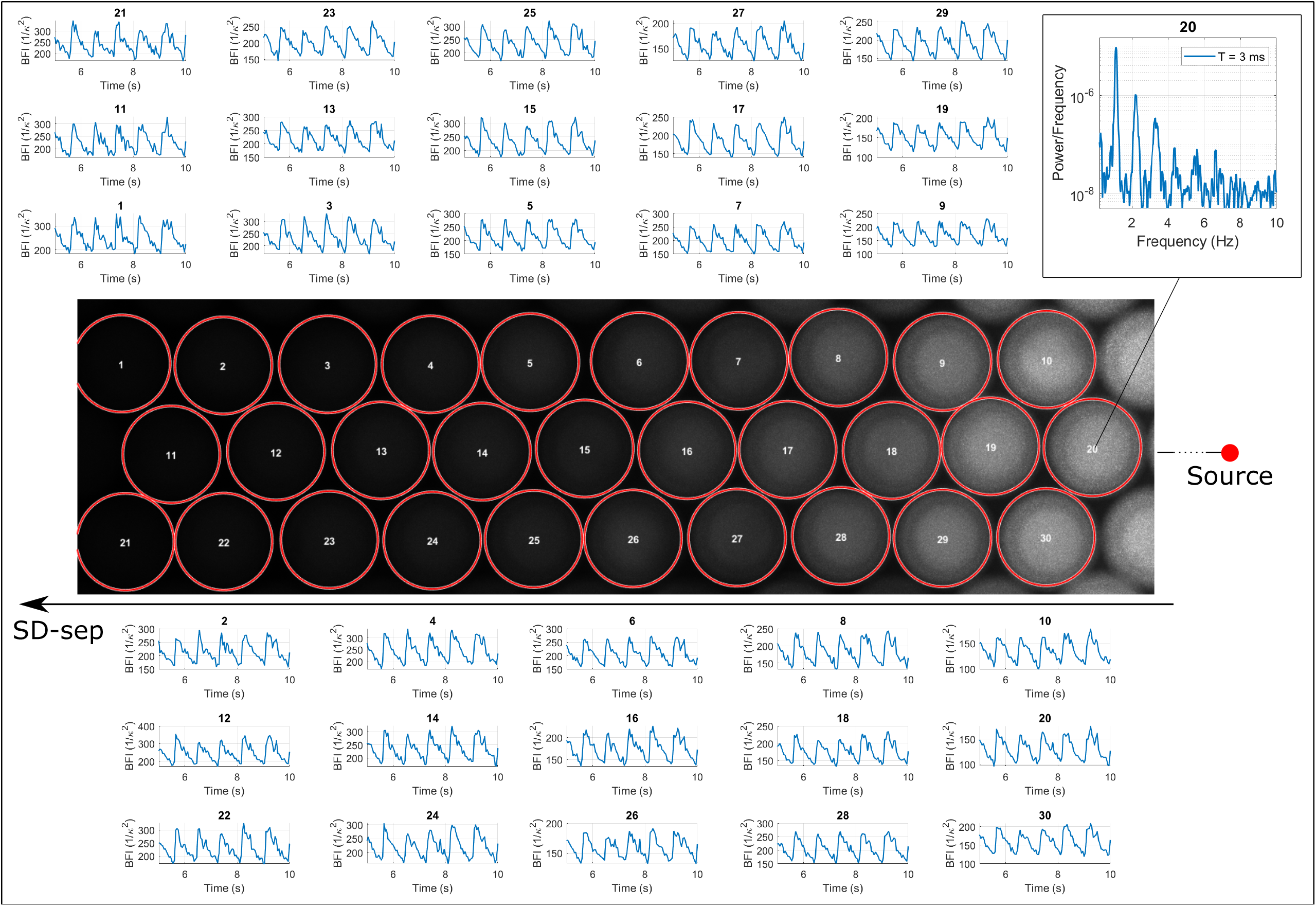
Examples of *BF I* pulsatile signal obtained from each lens of the array with an acquisition rate of 20 *Hz* and exposure time 3 *ms*. Top right: Fourier transform of the *BF I* obtained from lens #20. Source-detector separations range from 11.15 *mm* (lens #20) to 23.05 *mm* (lenses #1 and #21).

## 4. Discussion

In this study we have demonstrated the feasibility of a novel SCOS/SCOT approach based on a micro-objective array applied on a CMOS camera sensor. This approach allows direct skin contact measurements enabling high quality (∼ 10^4^ speckles per each lens field of view) simultaneous SCOS measurements at multiple source-detector separation with a single large area sensor, eliminating the need of fiber optics, which can introduce further noise to the measurements, being particularly sensitive to vibrations. In this respect, we note that while this demonstration utilized a fiber-coupled source, on-skin direct application of sources were previously demonstrated [11, 24, 25].

The use of a micro-objective array paves the way for additional customization, performance improvement and miniaturization of SCOS systems. We note that the camera used was *not selected* specifically for this purpose and hence forced the need for complex thermal management (i.e., bulk). Future implementations enabled by micro-optical technology could employ specialized camera designs without relying on bulky macro-optics, and could also integrate more compact cooling systems. The possibility of inserting custom filters between sensor and micro-objective array would also boost the performance and new applications of this modality. By using properly designed filters allowing different wavelength transmission for different micro-objective [16, 26], this approach will enable for example simultaneous multi wavelength measurements. By using a ND filter array with different transmittance at different source-detector separation, measurement dynamic range can also be easily improved, allowing to obtain an optimal signal simultaneously at different SDS.

The micro-objective array based SCOS system was then characterized with phantom and *in vivo* experiments. The static phantom measurements aimed to characterize basic properties of the speckle contrast images acquired. The average speckle size (∼ 2 pixels) and number (*>* 5000) within each lens field of view (∼ 17000 pixels) ensure adequate spatial sampling for statistical analysis, being comparable to many SCOS systems currently used [5, 9, 27, 28]. We note here that the analysis reported in this study was limited to a reduced region (approx. 60% radius) of each lens field of view, where lens transmission can be considered uniform. This region can easily be extended by advanced post-process analysis considering artifacts due to lens curvature and different lens transmission, further improving the SNR. Lastly, the intensity distributions from each lens field of view of dark signal and signal from the glass diffuser were calculated and reported in Figure 3 (bottom right panel), for an exemplary lens. While the dark distribution is symmetric (i.e., Gaussian), the intensity distribution from the static phantom is asymmetric, discriminating the speckle statistics from noise [9].

From the liquid phantom experiments, we were able to demonstrate the agreement of the measurements with the theory [1, 20]. As expected the decrease of the average intensity for increasing source-detector separation well agrees with the Beer-Lambert law, following an exponential decay. Additionally, the linear increase of the intensity Vs. camera exposure time confirms that the experiments in a region of camera linearity [6]. Lastly, the measured speckle contrast for varying source-detector separation and exposure time qualitatively matches theory expectations (equation 2).

Lastly, the *in vivo* test had the purpose of validating the capability of the system to track pulsatile blood flow. As highlighted in Figure 5, the quality of the pulses retrieved is good even at the largest source-detector separations measured, allowing the resolution of the complete cardiac cycle and the identification of the dicrotic notch [29]. This visual assessment was corroborated by Fourier analysis, revealing a SNR >250 for all configurations.

## 5. Conclusion

In this study we reported the first use of a SCOS system based on a CMOS camera based, micro-optical imaging approach for direct skin contact blood flow monitoring. By means of static and dynamic phantom measurements and *in vivo* tests we were able to characterize the system developed and demonstrate its suitability for being used for real time continuous monitoring of pulsatile blood flow. The performance of the system developed does not differ from many devices reported in literature during the last years, based on more traditional approaches as non-contact imaging systems or fiber optics based systems, while the use of a micro-objective array optimizes the detection area utilized by allowing simultaneous multiple source-detector separation measurements.

In conclusion, this study opens the path towards the development of miniaturized blood flow monitoring systems, embedding optical and electronic components directly into the skin contact probe.

## Footnotes

1 From hereon, we use SCOS to refer to both approaches.

